# Scalable gaussian process inference of neural responses to natural images

**DOI:** 10.1101/2023.01.13.523423

**Authors:** Matías A. Goldin, Samuele Virgili, Matthew Chalk

## Abstract

Predicting the responses of sensory neurons is a long-standing neuroscience goal. However, while there has been much progress in modeling neural responses to simple and/or artificial stimuli, predicting responses to natural stimuli remains an ongoing challenge. One the one hand, deep neural networks perform very well on certain data-sets, but can fail when data is limited. On the other hand, gaussian processes (GPs) perform well on limited data, but are generally poor at predicting responses to high-dimensional stimuli, such as natural images. Here we show how structured priors, e.g. for local and smooth receptive fields, can be used to scale up GPs to high-dimensional stimuli. We show that when we do this, a GP model largely outperforms a deep neural network trained to predict retinal responses to natural images, with largest differences observed when both models are trained on a very small data-set. Further, since GPs compute the uncertainty in their predictions, they are well-suited to closed-loop experiments, where stimuli are chosen actively so as to collect ‘informative’ neural data. We show how this can be done in practice on our retinal data-set, so as to: (i) efficiently learn a model of retinal responses to natural images, using little data, and (ii) rapidly distinguish between competing models (e.g. a linear vs a non-linear model). In the future, our approach could be applied to other low-level sensory areas, beyond the retina.

## 2 Introduction

Accurately predicting the responses of neurons in visual areas is an important goal in visual neuroscience. However, while there has been much progress in modelling sensory neural responses to simple and/or artificial stimuli (e.g. moving gratings/checkerboard) [1, 2], predicting neural responses to natural stimuli remains an ongoing challenge [3]. Even in the retina, arguably the simplest visual circuit, this problem is hard to solve [4, 5, 6].

One approach, that has had a great deal of success in recent years, is to train deep convolutional neural networks (CNNs) to predict neural responses to presented stimuli [7, 8]. However, this approach can suffer from two important limitations. First, CNNs typically require large amounts of data to train, whereas in many experimental settings data may be extremely limited. In such cases, CNNs will likely over-fit noise in the data, or alternatively, require strong regularisation which can bias the results [8]. Second, designing a deep CNN to work well on a given neural data-set may require a number of design choices (e.g. about the network architecture, non-linearities, etc.), and it may not be clear how each of these choices affect the final results [9].

Another approach has been to fit neural responses using a type of non-parametric Bayesian model, called gaussian processes (GPs) [10, 11, 12, 13]. These models can be trained on small datasets, in comparison to CNNs, while avoiding overfitting. They also estimate their own uncertainty, which can be useful for assessing the robustness of their predictions, as well as for applications such as active learning, where stimuli are chosen in closed-loop based on previously recorded responses [11, 12]. Unfortunately however, GPs typically perform very badly at predicting responses to highdimensional inputs, such as natural images or movies [14, 15]. As a result, in visual neuroscience, they have previously only been used to model neural responses to low-dimensional artificial stimuli, such as oriented gratings.

Here we propose a method to scale up GPs to predict neural responses to high-dimensional stimuli, such as natural images. To do this, we extend the approach of Park & Pillow [16], who showed how prior knowledge about visual neuron responses (e.g. that they have spatially local and smooth receptive fields), can be used to constrain and simplify inference of a simple, one-layer, linear-nonlinear (LN) model. We show that by incorporating this idea into a GP framework we can achieve good fits of retinal neuron responses to high-dimensional natural images, which out-perform the state-of-the-art CNN models, as well as the LN models considered by Park & Pillow and others. Moreover, our GP model continues to perform well even when trained on very little data, and when a CNN model breaks down. Finally, we describe how our GP framework can be used to design a closed-loop experiment, where one chooses which stimuli to present on-line, based on previously recorded responses. We demonstrate, using retinal recordings data, how our closed-loop algorithm can be used to quickly and efficiently learn a model of neural responses, as well as to discriminate between alternative competing scientific hypotheses.

## 3 Results

### 3.1 GP framework

Let us imagine we have a data-set, 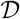, comprising *N* stimulus response pairs, (*x,r*). We denote the stimulus, *x*, by an *n_x_*-d vector (*n_x_* could be, e.g., the number of pixels in a presented image), while the response, *r*, corresponds to the number of spikes fired by a neuron in response to a presented stimulus. Our goal will be to use this data-set to learn a function, *f* (*x*) (which we call the neuron’s response curve) describing how many spikes the neuron fires on average to any given stimulus, *x*. Learning *f* (*x*) can be difficult due to three main factors: (i) if the stimulus is high dimensional (large *n_x_*); (ii) if we have limited, or noisy, data (small *N*); (iii) if the true response function, *f* (*x*), is highly non-linear.

We propose a gaussian-process (GP) framework to overcome these challenges. GPs are a type of non-parametric Bayesian model that return a posterior probability over response functions, given a training data-set of input-output pairs, (*x,r*) in our case. Our motivation for using this type of model is that, as with other Bayesian methods, they automatically apply Occam’s razor, so as to adjust the inferred model complexity based on the available data, and thus avoid over-fitting when *N* is small [17]. Further, they are non-parametric, and thus prevent us having to make strong assumptions about the form of the response function, *f* (*x*). Finally, as we will show later, the fact that GPs estimate the uncertainty in their predictions makes them useful for applications such as active sensing, where one chooses which stimulus to present in each trial in closed-loop [11, 12].

The output of a GP process is continuous and can take both positive or negative values. In contrast, spike counts are positive integers. Thus, to use GPs to tackle this problem, we assume that the GP output, λ(*x*), is transformed by a positive monotonic nonlinearity to generate the mean spike count, before applying Poisson noise to generate the observed spike count, *r* (**Fig. 1a**). Inference is performed using an approximate variational inference algorithm, in which we assume a multivariate gaussian posterior over λ, with parameters learned to maximise a lower bound on the log-likelihood with respect to the data. Further, we use the inducing point framework introduced by Hensman, Fusi & Laurence [18] (which involves conditioning the variational posterior over a limited number of *n* ≪ *N* ‘inducing’ data points) to make inference feasible when *N* is large (see Methods section 5.3).

**Figure 1:**
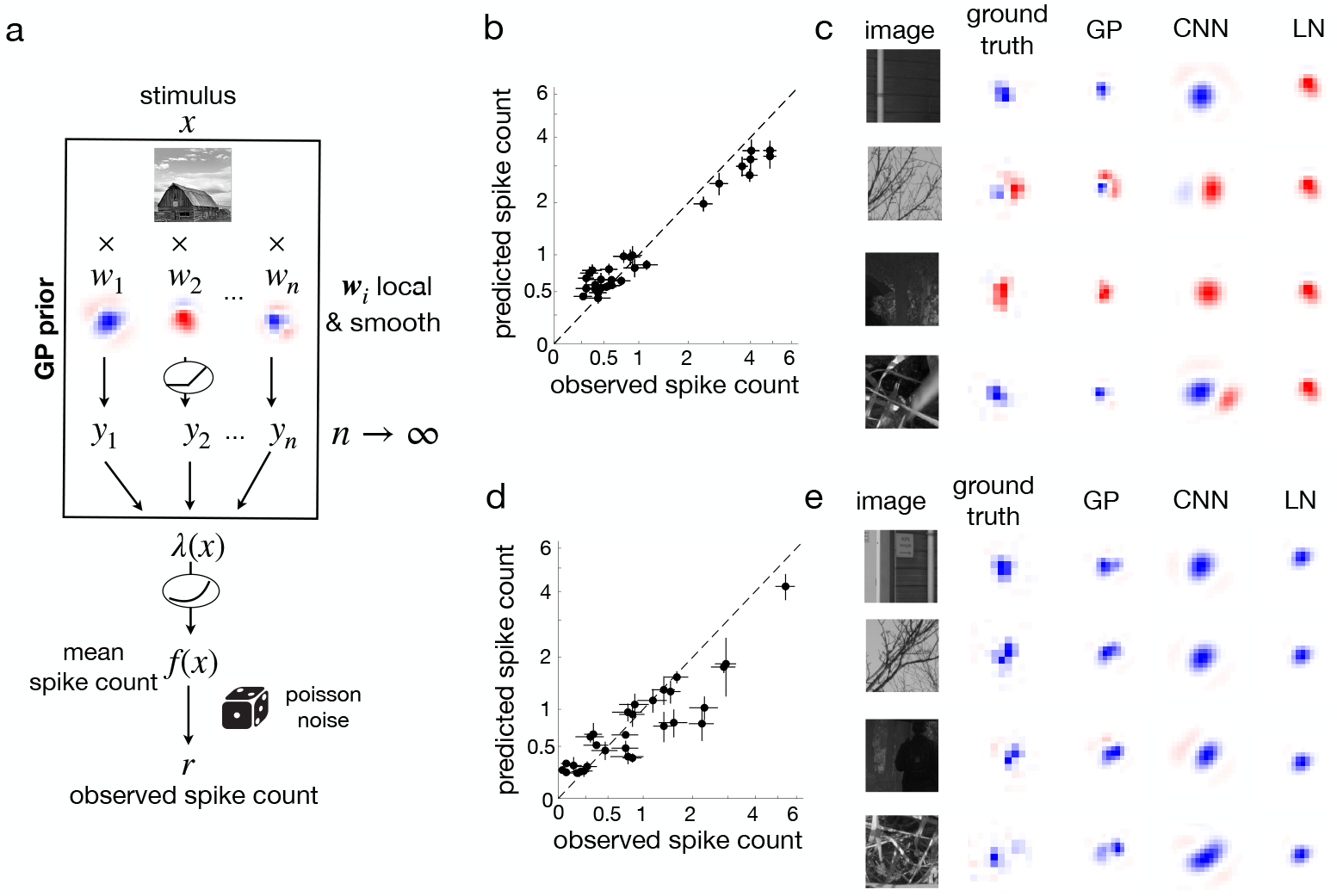
A novel GP model for neural responses to natural images. (a) Schematic of modelling framework. The stimulus (e.g. an image) is passed through a GP, then a positive non-linearity, before adding Poisson noise to obtain the observed spike count. Our proposed GP kernel was equivalent to using a 2-layer neural network, in which the first layer contains infinitely many spatial filters, w, followed by a positive threshold linear function. We assumed a strong prior for filters in the first layer to be spatially smooth and localised around the cell’s RF. (b) The predicted by the GP model versus the observed mean spike count for one retinal cell, for 30 repeated test stimuli. Vertical bars represent confidence intervals estimated by the model; horizontal bars represent confidence intervals estimated from data using bootstrap. (c) Examples of Local spike-triggered-average (LSTA) for this cell, estimated by a weighted average over a set of low amplitude noise patterns presented on top of a reference image (left). Both GP and CNN models were able to capture qualitative changes in the sign of the local STA, depending on the reference image. In contrast, an LN model could not capture stimulus-dependent changes in the local STA. (d)-(e) Same as (b)-(c) for another cell.

A GP is defined by its kernel function, *K* (*x, x′*), which specifies the prior covariance between the output function, λ(*x*), evaluated at *x* and *x*′ (i.e. cov (λ(*x*), λ (*x*′))). Different choices of kernel correspond to different prior assumptions about λ(*x*). A standard choice is the squared exponential kernel, which encapsulates our assumption that the function is smooth; it implies that similar inputs, *x* and *x*′, should generate similar outputs, λ(*x*) and λ(*x*′). Unfortunately however, this type of kernel doesn’t work well with high-dimensional inputs (i.e. large *n_x_*) [14]. This is because, in high-dimensions, the squared distance between any two stimuli in the training set, *x* and *x*′, is almost always very large. Thus it becomes hard, if not impossible, to predict the GP output to new high-d stimuli using just the assumption that the output function is smooth. We can partially overcome this problem by using ‘non-stationary’ kernels, which allow for long-range correlations between the response to different high-dimensional stimuli (at the expense of stronger assumptions about the response function). Here we choose to use an ‘arc-cosine’ kernel, proposed by [19] (Method section 5.1). This kernel is equivalent to assuming a two layer neural network, with infinitely many rectified-linear units in the middle layer (**Fig. 1a**).

Nonetheless, non-stationary kernels are still not sufficient to overcome the curse of dimensionality when nx is sufficiently large. In this case, we need additional assumptions to further simplify and constrain our inference. In the visual system, neurons typically respond to stimuli in a small region of space; i.e. they have spatially localised receptive fields (RFs). In addition, visual neuron RFs are typically spatially smooth. We can incorporate both these assumptions into our kernel, by assuming that the linear weights in the first layer of the two layer network are *a priori* spatially localised and smooth (Methods section 5.2). As is standard in GP inference, the hyper-parameters determining the location and smoothness of neural RFs are learned from data, so as to maximise the lower bound on the log-likelihood. Note that this extends the work of Park & Pillow [16] to a two-layer neural network, thus allowing us to consider non-linear response curves. While our focus here is the visual system, similar methods could be used for other sensory areas. For example, for audition we could use prior assumptions about spectro-temporal neural RFs.

### 3.2 Predicting retinal responses to natural images

To test the performance of our framework we trained it on a data-set comprising the recorded spiking responses of 41 retinal ganglion cells to a set of presented natural images (see Methods section 5.5 for experimental details [5]). This data-set gives rise to all of the challenges listed in the previous section since: (i) the stimulus is high dimensional (i.e. large number of pixels *n_x_* = 108^2^ = 11, 664); (ii) there are relatively few trials (*N* ≈ 3000 ≪ *n_x_*); (iii) from previous work, we know that retinal ganglion cell responses to naturalistic stimuli are often highly non-linear [5].

**Fig. 1b** & **d** shows the predicted versus the observed mean spike count for two example neurons, on 30 hold-out stimuli repeated 30 times each, that were not present in the training data-set. Since the model is probabilistic, it returns the uncertainty in its predictions (vertical lines), while the uncertainty in the data (horizontal lines) can be estimated using bootstrap. For both cells, the model achieves a reasonable prediction of the observed spike count, given the estimated experimental and modelling uncertainty (see later for quantitative assessment).

As well as fitting the mean spike count, we wanted to see if the GP model was able to predict other qualitative aspects of how the neural response curve depends on the presented image. To do this, we plotted the inferred gradient of the firing rate, ∇ *f* (*x*) |_*x*_0__, for a selection of ‘reference images’ *x*_0_ (**Fig. 1c** & **e**, left column). We compared this to a ‘ground truth’ estimate of the gradient, obtained by probing experimentally the neuron’s response to perturbations around the reference images, by presenting stimuli consisting of small white noise added on top of these images (**Fig. 1c** & **e**, second to left). For historical reasons, we call this estimated gradient, the ‘local spike-triggered-average’ (LSTA). It represents the the perturbation that produces the maximum change in the firing rate of the cell, given a reference image (Method section 5.5).

Most neurons exhibited an LSTA that varied qualitatively for different natural images, with occasional reversals in sign (**Fig. 1c**) and/or shape (**Fig. 1e**). In most cases, the GP model was able to reproduce these qualitative changes in the shape and sign of the recorded LSTA (**Fig. 1c** & **e**, middle column). In contrast, a single layer LN model was not able to reproduce these changes (**Fig. 1c** & **e**, right column), since by construction the gradient of the response function was the same for all stimuli (varying only in its total magnitude, due to the static non-linearity). We also compared our results to a 2-layer convolutional neural network (CNN), proposed previously to model this data-set [5] (**Fig. 1c** & **e**, second to right; see Methods section 5.6). While the CNN was able to produce most qualitative changes in the sign of the LSTAs, they appeared larger and less closely resembled the shape of the ground truth LSTAs, obtained using the perturbation stimuli. This could arise because of the restricted class of functions that can be captured by the CNN (i.e. a 2-layer network with a convolutional first layer), while in contrast, the GP can in theory fit any function, *f* (*x*), given enough data.

Next we wanted to evaluate quantitatively the performance of the model on this data-set. **Fig. 2a** shows the estimated lower bound on the log-likelihood achieved with the GP model (with arccosine kernel), compared with when we used a linear or quadratic kernel. Perhaps unsurprisingly, given the non-linear behaviour shown in **Fig. 1c** & **e**, the arc-cosine kernel greatly outperformed the linear kernel for every recorded cell. Interestingly, however, it also performed better than the quadratic kernel, which assumes that λ(*x*) is quadratic, and which has been used previously to model the responses of neurons in early visual areas [20].

**Figure 2:**
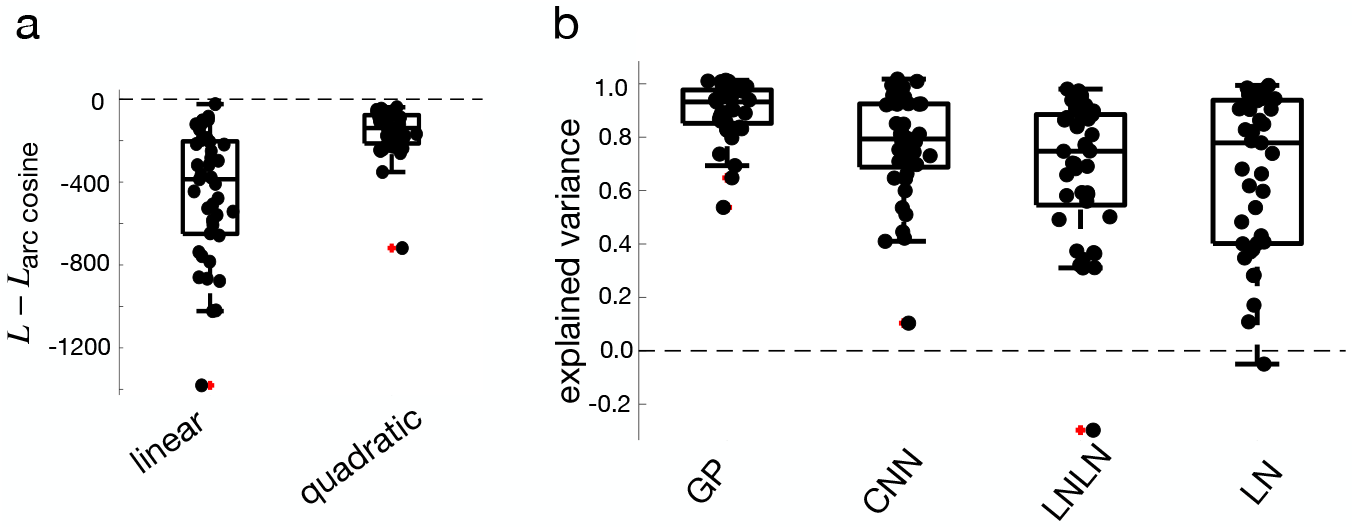
GP model outperforms other modeling approaches. (a) The estimated log-likelihood of the GP model for each cell (black dots), using either a linear or quadratic kernel, minus the loglikelihood achieved with the arc-cosine kernel. For every cell, the arc-cosine kernel performed best. (b) The explained variance for each cell on 30 repeated test stimuli for the GP model (with arc-cosine kernel), CNN model, LNLN model, and LN model. The GP model performed best, followed by the CNN.

To compare our model to other ‘non-Bayesian’ models, such as the two layer CNN plotted in **Fig. 1c** & **e** and a non-bayesian two layer LNLN model (Methods section 5.6), we plotted the noise corrected correlation, or ‘explained variance’, for each model for the 30 repeated hold-out images. Given the responses *r_i_*, we can divide them in even and odd trials and estimate their mean values, 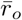 and 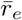. We defined the reliability as their correlation 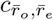. Then, we estimated the square noise-corrected correlation 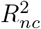 given a model prediction 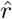:

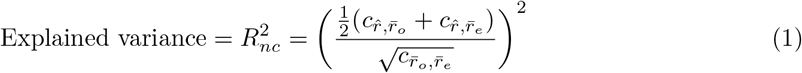

This metric, introduced by Keshishian et al. [21], corresponds to the *R*^2^ goodness of fit between observed and predicted firing rates, normalised based on the across-trial noise in the data. An explained variance of 1 means that the model predictions are as good as possible given the inherent noise in the data; an explained variance < 0 means that we would perform better by replacing the model predictions by a single number (the mean firing rate).

The GP performed well on this metric, with an explained variance >0.8 for 36/41 cells, and >0.6 for 40/41 cells. In contrast, while the CNN performed better than either the LNLN model and the LN model, it performed worse than the GP, with an explained variance >0.8 for only 19/41 cells, and >0.6 for 34/41 cells (**Fig. 2**).

### 3.3 Inference with little data

As stated earlier, one advantage of Bayesian models, such as GPs, is that they automatically apply Occam’s razor, so as to avoid overfitting when trained on a small number of trials. To test this, we trained our GP model on a small subset of the data, of *N* = 500 randomly selected trials. **Fig. 3a** shows the firing rate predicted by the GP model trained on 500 trials versus the model trained on *N* = 3160 trials, for the 30 repeated hold-out stimuli (here, we plot the cell shown in Fig. 1b-c). For this cell, the predicted firing rate is very similar in both cases, despite reducing the original data-set by a factor of 6. The same goes for the predicted LSTAs (**Fig. 3b**), which show very little difference when the model is trained on 500 versus 3160 trials. In contrast, the LSTAs predicted by the CNN appear drastically different when we reduce the training data-set.

**Figure 3:**
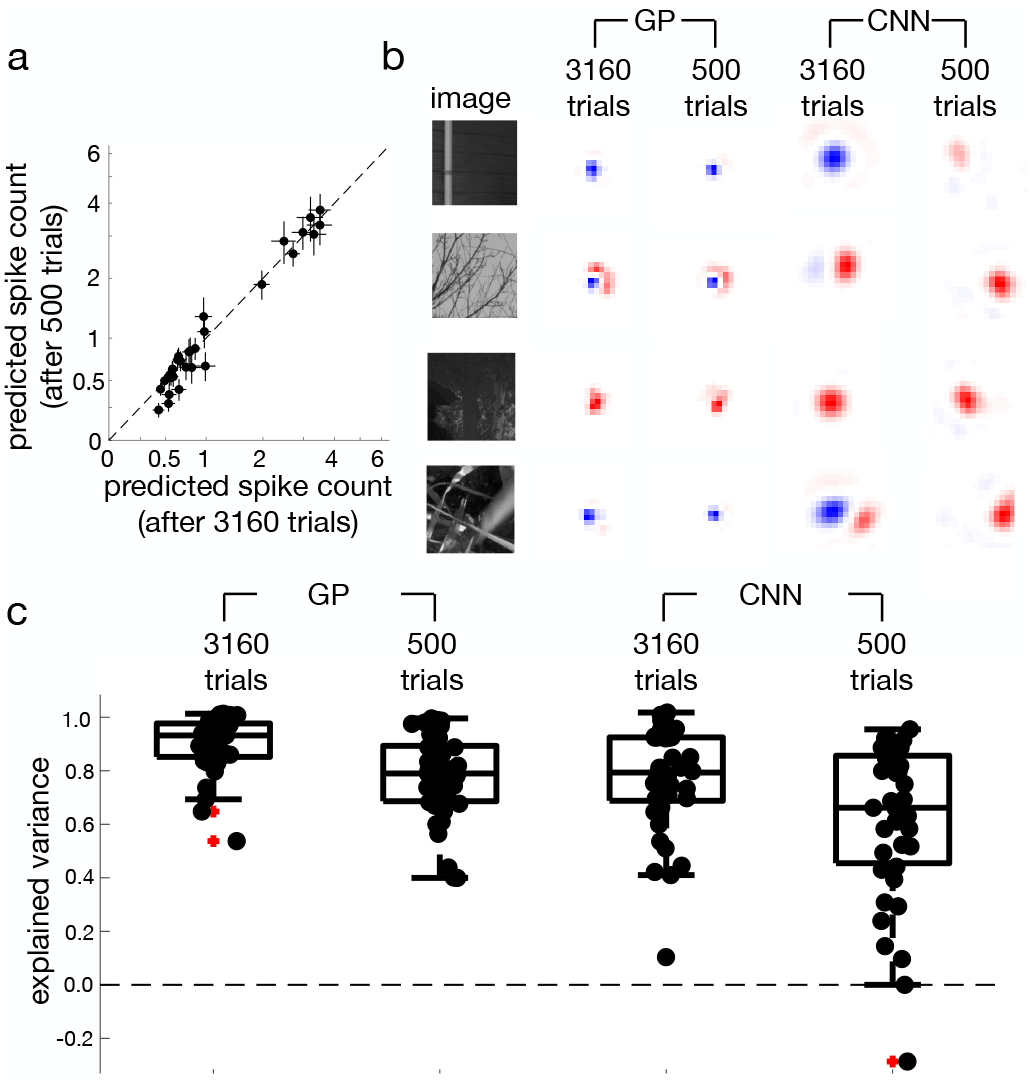
Performance of GP model when trained on little data. (a) Mean firing rate for one cell predicted by the GP model trained on 500 versus 3160 trials, on 30 test stimuli. (b) Estimated local STA predicted by the GP and CNN models when trained on 3160 vs 500 trials. For this cell, the local STAs predicted by the GP model are similar in each case, while the CNN predicts qualitatively different local STAs when trained on only 500 trials. (c) Explained variance achieved by the GP and CNN models when trained on 3160 vs 500 trials. While there is some decrease in performance of the GP when trained on fewer trials, its performance with 500 trials remains comparable to the CNN model trained on 3160 trials.

To see how the performance of both models reduces when we reduce the size of the training dataset, we plotted the explained variance for the 30 hold-out stimuli for both GP and CNN models. While the performance of the GP model was reduced when it was trained on only 500 trials, it was still comparable to the performance of the CNN trained on the full data-set of 3160 trials. In contrast, the explained variance achieved by the CNN model was greatly reduced when trained on fewer trials.

### 3.4 Active learning of the response curve

In the case where we have very little data, there may be very few stimuli that generate useful neural responses with which we can learn the neural response curve, *f*. To overcome this, we propose a closed-loop approach where at each instance of stimulus presentation, we use previously recorded neural responses to decide, or generate, which stimuli to present next. **Figure 4a** illustrates our proposed framework. After each recorded stimulus-response pair (*x, r*), the GP model is updated, resulting in new predictions for the mean and variance of the GP output, λ (*x*), to different natural stimuli (*μ* (*x*) & *σ* (*x*), respectively). We use these predictions to evaluate a utility function, *U* (*x*), which quantifies the ‘usefulness’ for learning the model of the neural response to a new stimulus, *x*. Finally, we choose the stimulus *x* for which *U* (*x*) is maximal to present on the next trial.

**Figure 4:**
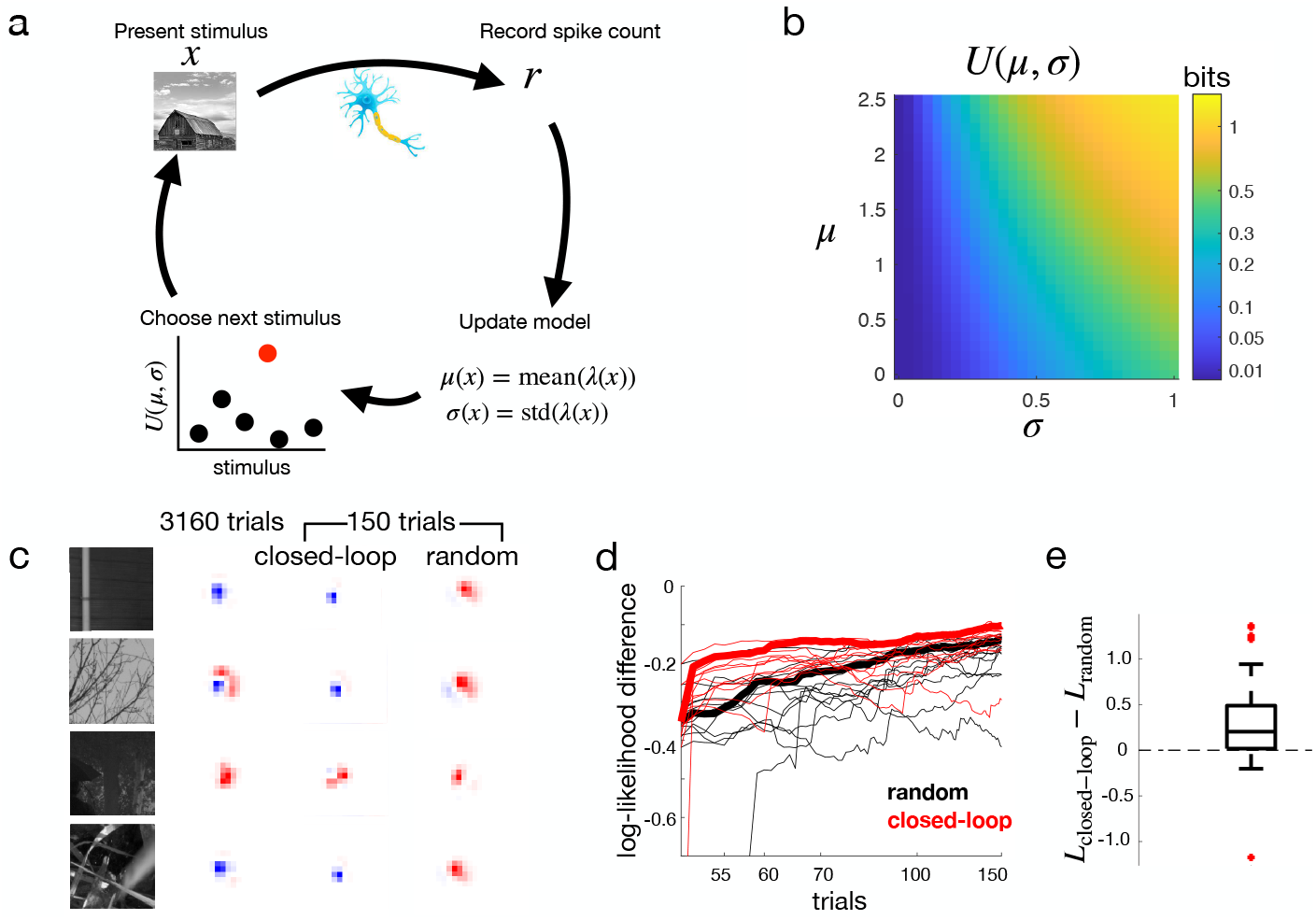
A closed-loop GP modeling framework. (a) Schematic of the procedure. At each instance, we use a recorded response to update the GP model, which predicts the mean and variance of the log firing rate, λ. We use this to compute a utility function, *U* (*x*), quantifying how informative each stimulus is about the response curve, *f*. (b) The utility function is largest for stimuli where the model predicts either high firing rates (large *μ*), or where predictions are uncertain (large *σ*). (c) Local STAs for one cell, estimated after 3160 trials (left), or after 150 stimuli, selected using the closed-loop algorithm (middle) or randomly (right). Local STAs more closely resemble the ground truth when we choose stimuli in closed-loop, rather than randomly. (d) The log-likelihood obtained with the GP model, evaluated on a test set of 1000 test stimuli, after training on a limited stimulus set chosen either in closed-loop (red) or randomly (black), minus the log-likelihood after training on the full data-set. Each line represents one run; thick lines represent the average over multiple runs. (e) The log-likelihood for all cells after a single 150 trial run, with stimuli chosen using the closed-loop algorithm, minus the log-likelihood when stimuli were selected randomly.

Our goal is for the recorded responses, *r*, to be maximally informative about the response function, *f*. Therefore, we define the utility function as the mutual information between the recorded response, *r*, and the response function, *f*, given a new presented stimulus, *x*, and data-set, 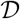:

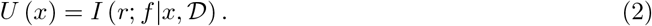

We can expand this as follows:

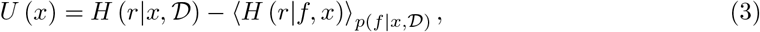

where the first term denotes the conditional response entropy, given *x* and 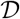, and the second term is the expectation (over 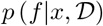) of the conditional response entropy, given *f* and *x*. Thus, we seek x for which the model is marginally most uncertain about *r* (high 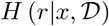), but for which individual settings of *f* are confident (low 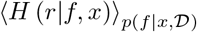).

The above utility function was introduced in the context of binary classification by Houlsby et al. [22]. In Methods section 5.4.1, we show how it can be well approximated for the Poisson observation model considered here, as a closed form expression of the mean and variance of λ (*x*) (*μ* (*x*) & *σ* (*x*)^2^, respectively). For a Poisson model, *U* increases monotonically with both *μ* and *σ* (**Fig. 4b**). Thus, *U* is largest when either: (1) the model predictions are uncertain (large *σ*), so that the first term of Eqn 3 is large; or (2) the predicted firing rate, and thus the signal-to-noise ratio, is large (large *μ*), so that the second term of Eqn 3 is small. (Note that, in contrast, in a standard GP model with constant noise *U* (*x*) doesn’t depend on the mean, *μ*.)

To work on-line, each step of the active learning algorithm must be executed quickly. In our work, the bottleneck was updating the hyperparameters of the GP, since both the approximate inference step (to update *μ* and *σ*) and utility function, *U*, could be computed quickly. To speed things up, therefore, we developed a ‘reduced’ GP model, for which most of the hyper-parameters were fixed, barring the multiplicative gain on the firing rate and the kernel magnitude, as these parameters could be updated quickly without altering the shape of the kernel *K* (*x, x*′). The parameters determining the location of each neuron’s RF were estimated using its responses to a flashed white noise stimulus, which was presented routinely at the start of each experiment. All other hyperparameters were kept the same for all neurons.

We used the data-set described above, comprising retinal responses to natural images, to show how our approach could work in principle. To mimic a closed-loop experiment, we initially only trained the model using a small subset of retinal responses to 50 natural images, chosen at random. We then revealed data points one at a time, either at random, or using the closed-loop algorithm described above.

**Fig. 4c** shows the learned LSTAs for one cell (the same cell shown in Fig. 1b-c) after 150 stimulus presentations, with stimulus-response pairs selected using the closed loop algorithm (middle), or at random (right). Here, learned LSTAs converged much more quickly to those inferred after 3160 trials (left) when we used the closed loop algorithm, versus choosing responses at random (right). To quantify the performance of the closed-loop algorithm, we measured the log-likelihood of the learned model, in predicting responses to a ‘test’ data-set (not used for training). Note that, while the closed-loop was performed using the ‘reduced’ GP model (with fixed hyperparameters) as described above, the plotted log-likelihood, used for evaluation, was computed using the full GP model, with all hyperparameters learned using the available stimulus response pairs at each step. As we can see in **Fig. 4d**, despite there being considerable variability across experimental runs, using a closed-loop algorithm to pick stimuli resulted in a consistently higher log-likelihood, compared to when stimuli are chosen at random. This was the case across the population of recorded cells, with a significantly higher log-likelihood for the closed-loop vs random stimulus selection, after performing one run of 150 trials on each cell (**Fig. 4e**).

### 3.5 Actively distinguishing between different models

Often, the goal is not to learn the response function, *f*, but rather to test whether a specific scientific hypothesis, *h*, is true (*h* = 1) or false (*h* = 0). We propose a closed-loop approach to address this problem using limited data. We adapt the framework described above by learning two different GP models in parallel, one for each experimental hypothesis (**Fig. 5a**). We then use these models to compute a utility function, *U* (*x*), which quantifies how much information the response to a given stimulus is expected to convey about the scientific hypotheses, *h*. Finally, we choose the stimulus that maximises *U* (*x*) to present on the next trial.

**Figure 5:**
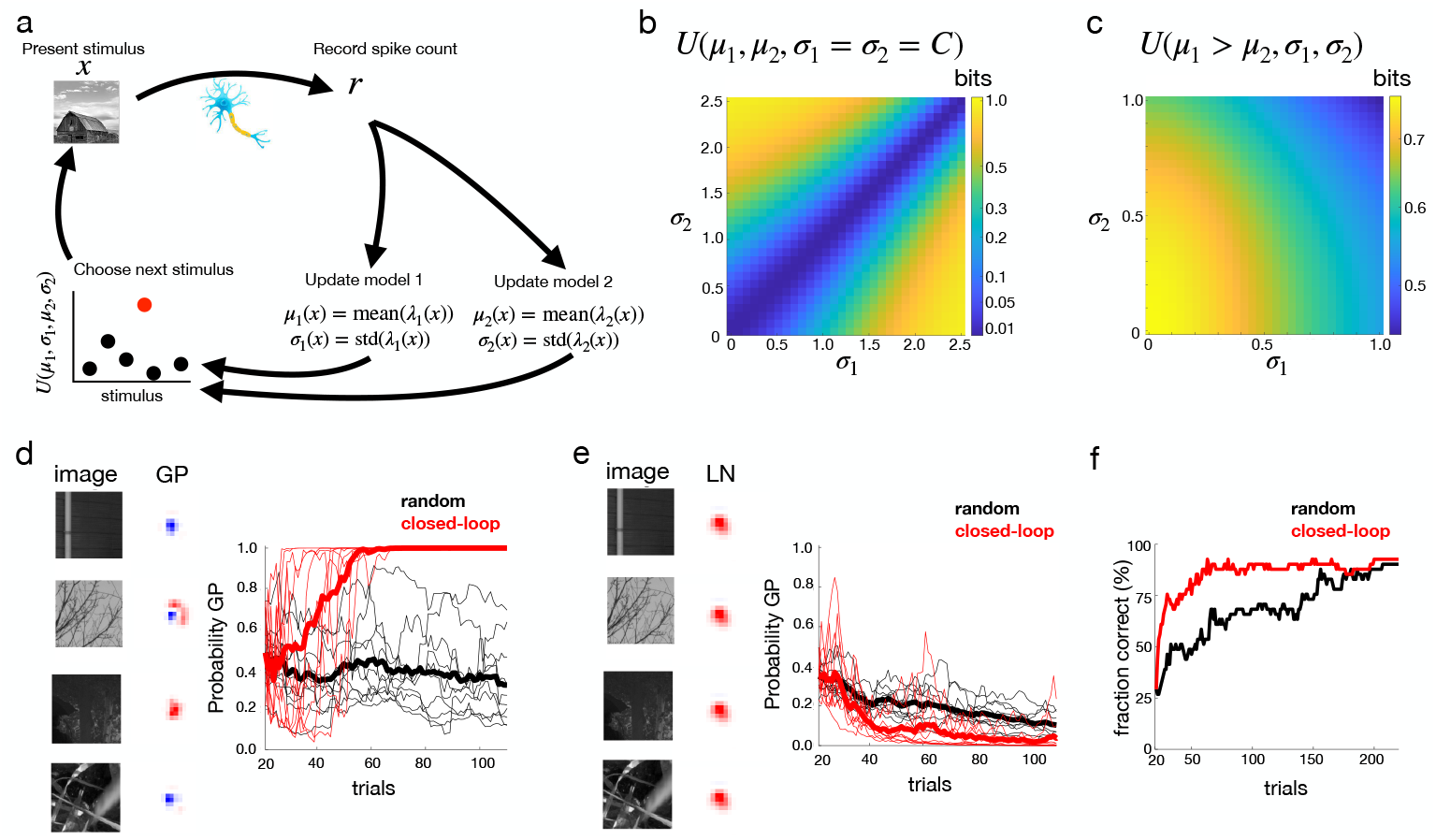
A closed-loop GP framework for distinguishing between models. (a) Schematic of the procedure. Here we trained two different, competing, models simultaneously. The utility function, U, quantifies how informative the responses to each stimulus are in distinguishing between these two models. (b) The utility function is largest when the two models predict very different mean firing rates (i.e. *μ*_1_ ≠ *μ*_2_). (c) The utility function decreases with the model uncertainty (i.e. increasing *σ*_1_ and/or *σ*_2_). (d) We generated synthetic data from a non-linear GP model, with local STAs shown here. We used the GP model to estimate the probability that synthetic data were generated from the non-linear vs a linear model. When stimuli were chosen in closed-loop, the algorithm was consistently quicker at correctly identifying the model class (red), compared to when stimuli were chosen randomly (black). (e) Same as panel d, but with synthetic data generated from a linear model, with local STAs shown on the left. (f) For each recorded cell, we used the GP algorithm to classify whether the data was better explained by a non-linear vs a linear model. Here we plot the fraction of cells where the model returns the ‘correct’ classification (i.e. the classification obtained after 3160 trials), when stimuli were selected in closed-loop (red) or randomly (black).

For the utility function, we choose the Shannon mutual information between the response, *r*, and hypothesis, *h*, conditioned on the stimulus *x*, and data-set, 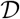:

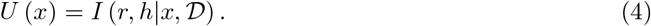

Expanding, we have:

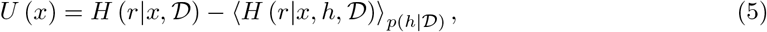

where the first term denotes the conditional response entropy, given *x* and 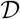, and the second term is the expected response entropy (over 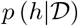), given hypothesis, *h*, and data *x* and data-set 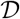.

In Methods section 5.4.2, we show how *U* (*x*) can be well approximated for the Poisson observation model considered here, as a closed form expression of the mean and variance of λ (*x*), given *h* = 0 and 1, (*μ*_1_ (*x*), *μ*_2_ (*x*), *σ*_1_ (*x*)^2^ & *σ*_2_ (*x*)^2^). *U* (*x*) is highest when both models give very different predictions about the mean firing rate, (i.e. when *μ*_1_ = *μ*_2_; **Fig. 5b**), but when the uncertainty associated with both models is small (i.e. *σ*_1_ and *σ*_2_ are small; **Fig. 5c**). Note that this contrasts with the utility function described in the previous section, for active learning of the response function, which increased monotonically with the uncertainty, *σ* (Fig. 4b).

To see how our approach could work in practice, we considered the case where the two hypothesis, to be tested are: (*h* = 0) neural responses are quasi-linear (i.e. they can be described by a GP with linear kernel, followed by static non-linearity); (*h* = 1) neural responses are non-linear (i.e. they can be described by a GP with arc-cosine kernel, followed by static non-linearity). We created a synthetic data-set, generated using a GP model (fitted to a neuron in the real data-set) with either a linear (*h* = 0) or non-linear kernel (*h* = 1) (**Fig. 5d-e**, left). To test our approach, we used our GP framework to infer the probability that *h* = 1, after a small number of trials with the synthetic data-set. We compared the case where stimuli were chosen using the closed-loop algorithm, or at random. As intended (**Fig. 5d-e**, right), we were significantly faster at correctly identifying whether the synthetic data was generated using a linear or a non-linear model when stimuli were chosen using the closed-loop algorithm, and not at random.

Finally, we sought to evaluate the performance of the closed-loop algorithm on the real data-set. As here there was no ground truth available, we defined the ‘true’ model class for each neuron to be the preferred model (i.e. linear or non-linear) selected by the reduced GP algorithm when trained on the full data-set of 3160 trials. We then plotted how quickly the algorithm converged to ‘correctly’ identify the class of each cell versus the number of presented stimulus-response pairs, with stimuli selected either using the closed-loop algorithm or at random (**Fig. 5f**). As hoped, the algorithm was significantly quicker at correctly classifying the model class when stimuli were selected using the closed-loop algorithm, and not at random.

## 4 Discussion

Previous work using GP models to fit neural responses was restricted to low-dimensional, artificial stimuli, such as moving oriented gratings [11, 12, 13]. This is likely because most commonly used stationary kernels (e.g. the squared-exponential or Matern kernel) rely on simple assumptions about the response function (e.g. that it is smooth), which don’t scale well to high-dimensional inputs such as natural images [14, 15]. To scale to higher input dimensions, additional assumptions are required, such as the fact that neural RFs are spatially local and smooth [16, 23, 24], which act to reduce the effective dimensionality of the input. As we showed, this works well for modelling retinal neurons, and is also expected to work for other low-level visual areas such as V1 and V2, where receptive fields of neurons cover restricted areas of the visual scene. In addition, our approach could be adapted to other sensory areas, such as the primary auditory cortex, where neurons respond selectively to a small range of spectral frequencies.

To demonstrate the advantages of GP processes to model neural data, we started by considering neural responses to a static image. In the future it would be interesting to extend our model to neural responses to dynamic stimuli, such as natural movies [25]. This would of course increase the dimensionality of the input, and thus the difficulty of inference. Nonetheless, additional prior assumptions, such as local and smooth temporal response functions [24], could be used to keep inference tractable.

We fitted the GP model on each neuron separately, in contrast to the CNN model, whose first layer was trained using data from the entire recorded neural population [5]. Despite this, the GP model outperformed the CNN model (**Fig 2b**), with strongest differences observed when both models were trained on very little data (**Fig. 3b-c**). This is likely because Bayesian models, such as GPs, adjust their complexity to avoid over-fitting when trained on limited data. In the future, the performance of the GP model could likely be increased further by fitting it jointly on multiple recorded neurons, as we did for the CNN. While a GP model doesn’t contain multiple layers, one way to do this would be to assume that the responses of different neurons are obtained by linearly combining a small number of latent GPs [26, 27, 28] (while performing a spatial translation on the input, to take into account the different RF locations). For the retina, these latent GPs could have a biological relevance, as corresponding to different cell-types [4, 29]. Standard methods for classifying retinal cells types functionally are based on the response to simple stimuli (full field luminance amplitude or frequency variations, moving bars). However, the extension of our methods using latent GPs could classify the cells using more relevant stimuli (natural images), with the addition of providing a model for each cell [30].

We found that best results were obtained with a non-stationary arc-cosine kernel, proposed by Cho & Saul [19]. One reason could be that this kernel corresponds to a 2-layer neural network, which roughly approximates the retinal architecture (where there is an intermediate bipoloar cell layer, before the recorded ganglion cells). In their original paper [19], Cho & Saul, showed how their kernel could be simply extended to incorporate more neural layers. While we found this did not significantly improve performance for the retinal data-set, it is possible that this could help for higher-level visual areas beyond the retina, such as V1 and V2. Indeed, to model these higher level areas, future work could look at extending the kernel used here, to include aspects of convolutional [31] and/or deep [32] neural networks.

In recent closed-loop experiments using deep neural networks, stimuli were chosen so as to elicit a maximal response in recorded neurons [33, 34, 35]. In contrast, our approach looks for stimuli that are most informative about the entire response curve, *f* (*x*), and not just its maximum, maxx *f* (*x*). This can be seen in the acquisition function, *U* (*x*), used to choose new stimuli (Fig. 4b), which favoured stimuli for which there was high uncertainty (large *σ*), as well as a high predicted firing rate (large *μ*). Note that it would not be possible to do this for standard deep neural networks, as they do not compute the uncertainty in their predictions (*σ*). Moreover, our approach can be extended to choose stimuli, in closed-loop, that allow us to best address the specific scientific question at hand [36] (Fig. 5). To illustrate this, we considered a simple question: are retinal neurons best described with an LN or and LNLN model? Future work could consider a range of different questions. For example, we could choose stimuli such that we can best ascertain which groups of neurons in the retina belong to the same cell-type [29, 30]. Alternatively, in higher level visual areas we could select stimuli so as to best test the effects of top-down processes such as visual attention and expectations.

## 5 Methods

### 5.1 Gaussian process model

We construct a two layer linear-nonlinear (LNLN) model of neural responses. The spike count, r is assumed to be drawn from a Poisson distribution with mean *f* (*x*) = *e*^λ(*x*)+λ_0_^, where *x* is an *n_x_*-d stimulus vector, λ_0_ is a constant bias term, and λ (*x*), is given by a weighted sum of *J* units:

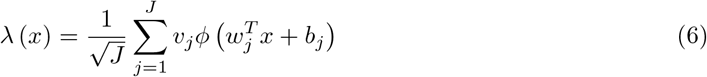

where *ϕ* is a non-linear basis function, *v_j_* are scalar weights, *w_j_* is an *n_x_*-d vector of linear weights, and *b_j_* is a scalar bias term.

We assume *a priori* that parameters, *w_j_, v_j_* and *b_j_*, are sampled from independent and identical distributions for different *j*. In this case, under fairly general conditions, in the limit *J* → ∞, we can use the central limit theorem to see that λ (*x*) is a Gaussian process, with mean, *μ*_0_ ≡ 〈λ (*x*)〉, and covariance, *K* (*x, x*′) ≡ cov (λ (*x*), λ (*x*′)).

With no loss of generality, we assume that weights in the second-layer are sampled from a standard normal distribution, 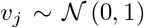. We assume a multivariate gaussian prior over input weights in the first layer, with zero mean and covariance *C*, while the bias term, *b_j_* is drawn from a gaussian with zero mean and variance 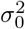. It is straightforward to show that in the limit *J* → ∞, the prior mean of λ is zero, while its covariance is given by:

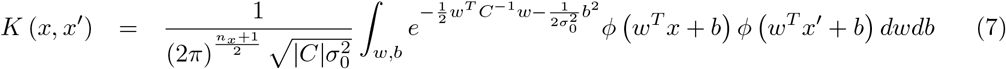

The above kernel depends on the form of the non-linearity, *ϕ*. Here we consider a rectified-nonlinearity, *ϕ*(*x*) = max (*x*, 0), which gives the following kernel:

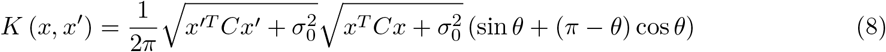

where,

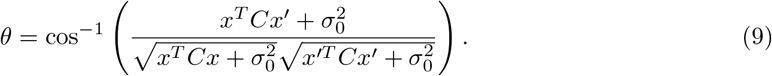

In the special case where *C* = *I* and *σ*_0_ = 0 this proposed kernel becomes equivalent to the arc-cosine kernel proposed by [19].

As a control, we also considered the gaussian kernel,

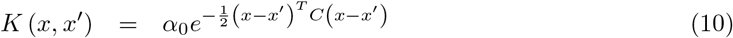

where *α*_0_ is a positive constant, as well as a linear kernel:

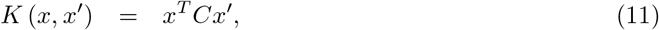

and a quadratic kernel:

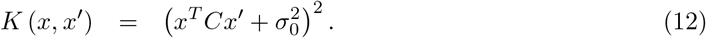

### 5.2 Spatially local and smooth receptive fields

Extending the work of Park & Pillow [16], we assume a prior distribution over the weights in the first layer, 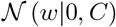, to enforce spatially local, smooth, receptive fields. To do this, we associate each element, *w_i_*, with a spatial location, *ξ_i_*. We then construct a covariance matrix with elements:

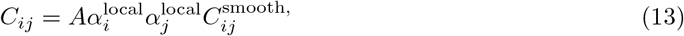

where *A* is a positive constant. *α*^local^ enforces *w* to be localised around a spatial location *ξ*_0_, and to be zero everywhere else. It is given by,

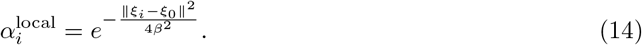

*C*^smooth^, enforces *w* to be spatially smooth, and is given by,

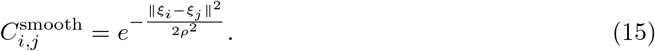

The covariance function, *C* contains 5 hyperparameters. *β* and *ρ* determine how spatially localized and smooth the receptive field is, respectively. A 2-d coordinate *ξ*_0_ determines the spatial location of the receptive field. Finally, a scalar, *A*, determines magnitude of *w*.

### 5.3 Inference

We seek to learn the posterior distribution, 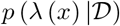, where λ (*x*) is the log-firing rate, and 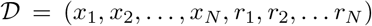, is the observed data-set. Unfortunately, exact inference with a GP model quickly becomes infeasible as the number of data points, *N*, becomes large. To get around this, we define a variational distribution, 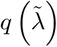, over a set of *n* ≪ *N* ‘inducing-points’, 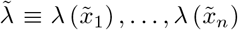. Predictions are then made by computing 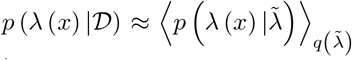. Doing this reduces the cost of inference from *O* (*N*^3^) to *O* (*n*^2^*N*). In the experiments described here we chose *n* = 250, since little improvement was observed when n was increased further. For the closed-loop experiments we set *n* = *N*.

We choose *q* to be a multivariate gaussian with mean m and covariance *V*, 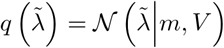. We optimise *m* and *V* by maximising the following lower bound on the log-likelihood:

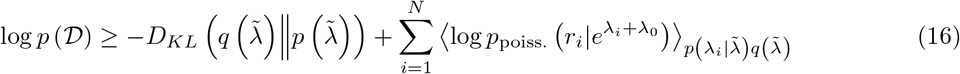

where *D_KL_* (*p*||*q*) is the Kullback-Leibler divergence between *p* and *q, p*_poiss_. is a Poisson distribution, and we have defined λ_*i*_ ≡ λ (*x_i_*).

The KL-divergence divergence term takes the following form:

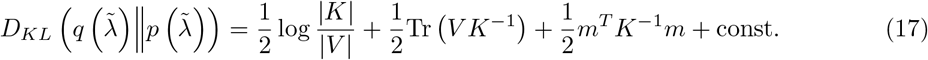

where *K* is an *n* × *n* kernel-gram matrix with elements, 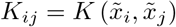.

The second term is the average log-likelihood of the data,

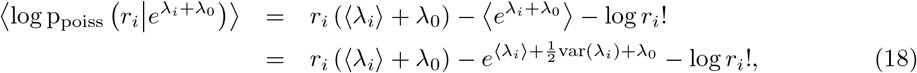

where,

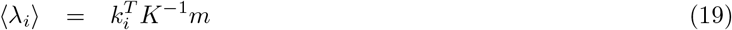

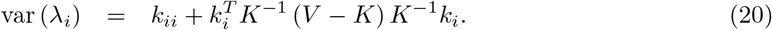

Here, *k_ii_* is a scalar given by *K*(*x_i_, x_i_*), while *k_i_* is an *n*-d vector, with *j^th^* element 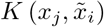.

The lower bound on the log-likelihood, in Eqn 16, is concave with respect to *m* and *V*. We can efficiently maximise this lower bound by performing the following Newton updates:

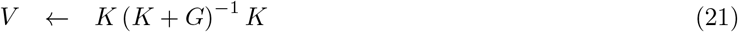

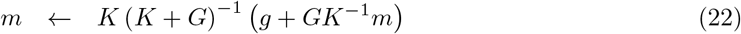

where:

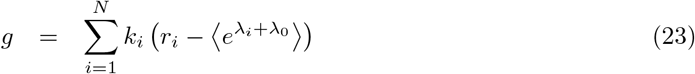

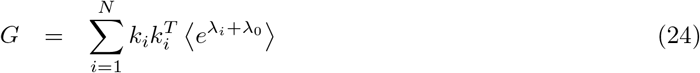

The hyperparameters determining the kernel matrix, *K*, and bias term, λ_0_, are then updated using gradient-descent to maximise the lower-bound on the log-likelihood, for fixed *m* and *V*. Optimisation proceeds via an EM algorithm, by alternately updating 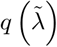 for fixed *θ* and λ_0_ (the E-step), then updating *θ* and λ_0_ for fixed 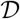 (the M-step).

After learning, the mean and variance of the predicted firing rate are given by:

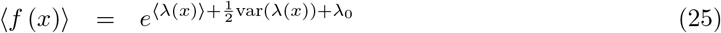

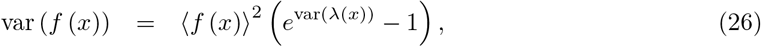

where we recall that 〈λ_*i*_〉 & var (λ_*i*_) are given by Eqns 19 & 20, respectively.

### 5.4 Optimal experimental design

#### 5.4.1 Active learning of the response function

We seek to choose a new stimulus so as to maximise the mutual information between the response, *r*, and the firing rate, *f*, given data, 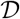 and a new stimulus, *x*:

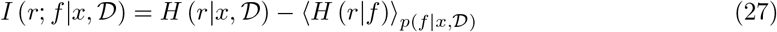

The first, response entropy, term is given by:

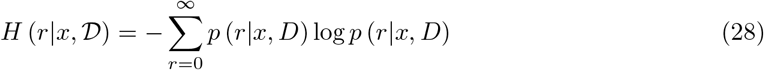

where,

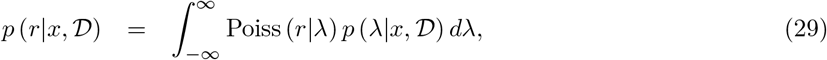

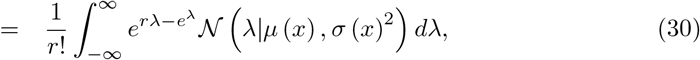

and *μ* (*x*) and *σ* (*x*)^2^ are defined here as the posterior mean and variance of λ (*x*) given previous observations 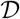. From here-in we will drop the dependence on *x* for notational simplicity. While the above integral is intractable, we can closely approximate it using Laplace approximation:

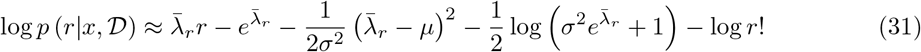

where

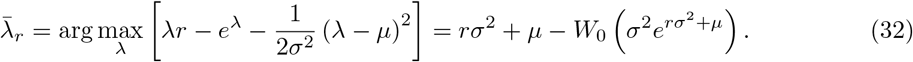

where *W*_0_ is the zeroth branch of the Lambert W-function. We then substitute this approximation into Eqn 28 to find the entropy. Note that while in theory, the sum in Eqn 28 goes to infinity, in practice we can do fairly well with a truncated version of this sum, so long as the mean firing rates are small (which is the case for our data-sets).

Next, we need to find the conditional noise entropy, given by:

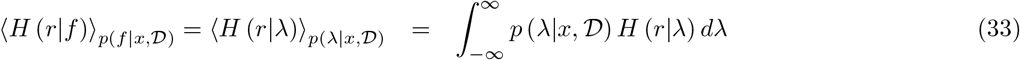

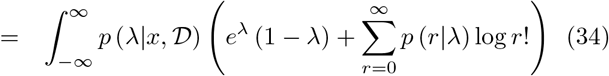

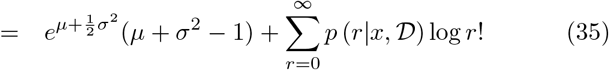

The last term can be approximated using Eqn 31. Again, the summation can be truncated with little loss in accuracy if firing rates are small.

#### 5.4.2 Active learning of the model class

Here, we seek to maximise the mutual information between the response, *r*, and the model class, *h*, given data, 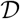 and new stimulus, *x*:

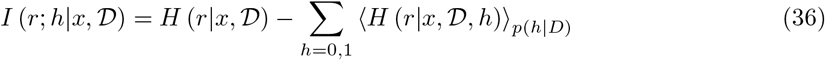

The first term is the entropy of the response if we don’t know the model class. To obtain this, we need to compute the posterior distribution of responses, if the model class is unknown, given by:

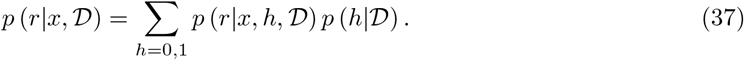

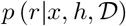 is approximated using the Laplace approximation described in the previous section (Eqn 31). 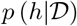 is given by:

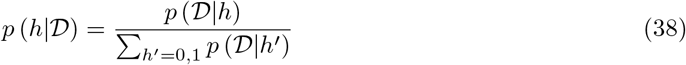

where 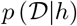 is approximated using our variational lower bound (Eqn 16), which we compute at each stage of inference.

### 5.5 Experimental details

We used a data-set consisting of the responses of 41 retinal ganglion cells of a mouse to various visual stimuli, reported in Goldin et al., 2022 [5]. In this experiment, perturbed and unperturbed natural images, together with pure white noise stimuli, were presented to the retina, while recording extra-cellularly retinal ganglion cells with a multi-electrode-array. There were 3190 unperturbed natural images, 30 of which were repeated 30 times to form the test set, and 8000 perturbed images (i.e. natural image plus white noise perturbation). Perturbed and unperturbed natural images were grey-scale (8-bits) and 864×864 pixels, which we down-sampled to 108 × 108 for modelling.

Characterization of the receptive field (RF) size and position of cells was done with white noise random checkerboard stimuli at 30 Hz. Receptive fields were estimated using a classical ‘Spike Triggered Average’ (STA), and were fitted with a two dimensional Gaussian function to estimate its parameters.

A ’local spike-triggered-average’ (LSTA) of each cell (**Fig. 1c** & **e**) was calculated for each reference image by counting the spikes evoked between 30 and 350 ms after each perturbed natural image presentation, and averaging the perturbation patterns weighted by their corresponding spike counts.

Spike count responses *r* evoked by a cell to unperturbed natural images, and used for model fitting were also obtained by counting the number of spikes between 30 and 350 ms after each natural image presentation.

### 5.6 CNN model

The CNN model is described in Goldin et al. 2022 [5]. It consists of a two layer neural network. The first layer was convolutional, with four two-dimensional kernels, learned from data, and common to all the recorded cells. The second layer consisted of one filter for each cell, followed by a nonlinearity. To reduce dimensionality, we factorized the weights of each filter into a two-dimensional spatial mask and a feature weight vector. In the first layer, we used Laplacian regularization on the convolutional kernels, and on the second layer, we used *L*_1_ regularisation on the spatial weights and on the feature weights. Regularisation hyperparameters were selected using cross-validation. This was done separately when the model was trained on a reduced data-set (**Fig 3**).

### 5.7 LNLN model

As for the CNN, the LNLN consists of a two layer neural network, where each layer linearly combines its inputs before applying a (soft-plus) non-linearity (with bias and gain learned from data). In contrast to the CNN, there are no shared weights between neurons: each neuron was trained separately. The first layer consists of two, 7 × 7 convolutional kernels, 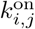 and 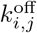 (where *i* and *j* index space) that compute two convolutions of the image, followed by a soft-plus nonlinearity. The 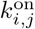 and 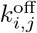 were chosen to be 2D Gaussian spatial profiles of opposite polarity but sharing the same spatial scale (following previous results [5]), and were kept constant during training. The second layer consists of linear filter weights 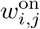 and 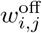, followed by a soft-plus non-linearity, which is applied to the output of the first layer. We assume a Poisson observation model for training. We used a *L*_1_ and a Laplacian regularization to train the linear filters of the second layer, whose hyperparameters were selected by cross-validation.

## Notes

### Competing Interest Statement

The authors have declared no competing interest.

## References

[1] Pillow, J. W., Shlens, J., Paninski, L., Sher, A., Litke, A. M., Chichilnisky, E. J., & Simoncelli, E. P. (2008). Spatio-temporal correlations and visual signalling in a complete neuronal population. Nature, 454(7207), 995–999.

[2] Bartsch, F., Cumming, B. G., & Butts, D. A. (2022). Model-based characterization of the selectivity of neurons in primary visual cortex. Journal of Neurophysiology, 128(2), 350–363.

[3] Willeke, K. F., Fahey, P. G., Bashiri, M., Pede, L., Burg, M. F., Blessing, C., … & Sinz, F. H. (2022). The Sensorium competition on predicting large-scale mouse primary visual cortex activity. arXiv preprint arXiv:2206.08666.

[4] Klindt, D., Ecker, A. S., Euler, T., & Bethge, M. (2017). Neural system identification for large populations separating “what” and “where”. Advances in Neural Information Processing Systems, 30.

[5] Goldin, M.A., Lefebvre, B., Virgili, S., Pham Van Cang, M.K., Ecker, A., Mora, T., Ferrari, U., & Marre, O. (2022). Context-dependent selectivity to natural images in the retina. Nature Communications, 13(1), pp.1–12.

[6] Maheswaranathan, N., Kastner, D. B., Baccus, S. A., & Ganguli, S. (2018). Inferring hidden structure in multilayered neural circuits. PLoS computational biology, 14(8), e1006291.

[7] Kriegeskorte, N. (2015). Deep neural networks: a new framework for modelling biological vision and brain information processing. Annu. Rev. Vis. Sci 1 (2015): 417–46.

[8] Yamins, D. L., & DiCarlo, J. J. (2016). Using goal-driven deep learning models to understand sensory cortex. Nature neuroscience, 19(3), 356–365. Chicago

[9] Tanaka, H., Nayebi, A., Maheswaranathan, N., McIntosh, L., Baccus, S., & Ganguli, S. (2019). From deep learning to mechanistic understanding in neuroscience: the structure of retinal prediction. Advances in neural information processing systems, 32.

[10] Williams, C. K., & Rasmussen, C. E. (2006). Gaussian processes for machine learning (Vol. 2, No. 3, p. 4). Cambridge, MA: MIT press.

[11] Park, M., Horwitz, G., & Pillow, J. (2011). Active learning of neural response functions with Gaussian processes. Advances in neural information processing systems, 24.

[12] Park, M., Weller, J. P., Horwitz, G. D., & Pillow, J. W. (2014). Bayesian active learning of neural firing rate maps with transformed gaussian process priors. Neural computation, 26(8), 1519–1541.

[13] Greenidge, C. D., Scholl, B., Yates, J. L., & Pillow, J. W. (2022). Efficient decoding of large-scale neural population responses with Gaussian-process multiclass regression. bioRxiv, 2021–08.

[14] Binois, M., & Wycoff, N. (2022). A survey on high-dimensional Gaussian process modeling with application to Bayesian optimization. ACM Transactions on Evolutionary Learning and Optimization, 2(2), 1–26.

[15] Delbridge, I., Bindel, D., & Wilson, A. G. (2020). Randomly projected additive Gaussian processes for regression. In International Conference on Machine Learning (pp. 2453–2463). PMLR.

[16] Park, M., & Pillow, J. W. (2011). Receptive field inference with localized priors. PLoS computational biology, 7(10), e1002219.

[17] MacKay, D. J. (2003). Information theory, inference and learning algorithms. Cambridge university press, Chapter 28, page 343

[18] Hensman, J., Fusi, N., & Lawrence, N. D. (2013). Gaussian processes for big data. arXiv preprint arXiv:1309.6835.

[19] Cho, Y., & Saul, L. (2009). Kernel methods for deep learning. Advances in neural information processing systems, 22.

[20] Rajan, K., Marre, O., & Tkačik, G. (2013). Learning quadratic receptive fields from neural responses to natural stimuli. Neural computation, 25(7), 1661–1692.

[21] Keshishian M, Akbari H, Khalighinejad B, Herrero JL. Estimating and interpreting nonlinear receptive field of sensory neural responses with deep neural network models. 2020:1–24.

[22] Houlsby, N., Huszár, F., Ghahramani, Z., & Lengyel, M. (2011). Bayesian active learning for classification and preference learning. arXiv preprint arXiv:1112.5745.

[23] Huang, Z., Ran, Y., Oesterle, J., Euler, T., & Berens, P. (2021). Estimating smooth and sparse neural receptive fields with a flexible spline basis. arXiv preprint arXiv:2108.07537.

[24] Duncker, L., Ruda, K. M., Field, G. D., & Pillow, J. W. (2022). Scalable variational inference for low-rank spatio-temporal receptive fields. bioRxiv.

[25] Hoefling, L., Szatko, K. P., Behrens, C., Qiu, Y., Klindt, D. A., Jessen, Z., … & Euler, T. (2022). A chromatic feature detector in the retina signals visual context changes. bioRxiv.

[26] Yu, B. M., Cunningham, J. P., Santhanam, G., Ryu, S., Shenoy, K. V., & Sahani, M. (2008). Gaussian-process factor analysis for low-dimensional single-trial analysis of neural population activity. Advances in neural information processing systems, 21.

[27] Rutten, V., Bernacchia, A., Sahani, M., & Hennequin, G. (2020). Non-reversible Gaussian processes for identifying latent dynamical structure in neural data. Advances in neural information processing systems, 33, 9622–9632.

[28] Wu, A., Roy, N.A., Keeley, S., & Pillow, J.W. (2017). Gaussian process based nonlinear latent structure discovery in multivariate spike train data. Advances in neural information processing systems, 30.

[29] Baden, T., Berens, P., Franke, K., Román Rosón, M., Bethge, M., & Euler, T. (2016). The functional diversity of retinal ganglion cells in the mouse. Nature, 529(7586), 345–350.

[30] Trapani, F., Spampinato, G., Yger, P., & Marre, O. (2022). Differences in non-linearities determine retinal cell types. bioRxiv.

[31] Van der Wilk, M., Rasmussen, C. E., & Hensman, J. (2017). Convolutional gaussian processes. Advances in Neural Information Processing Systems, 30.

[32] Blomqvist, K., Kaski, S., & Heinonen, M. (2020). Deep convolutional Gaussian processes. In Joint European Conference on Machine Learning and Knowledge Discovery in Databases (pp. 582–597). Springer, Cham.

[33] Cadena, Santiago A., George H. Denfield, Edgar Y. Walker, Leon A. Gatys, Andreas S. Tolias, Matthias Bethge, & Alexander S. Ecker. ”Deep convolutional models improve predictions of macaque V1 responses to natural images.” PLoS computational biology 15, no. 4 (2019): e1006897.

[34] Ponce, C. R., Xiao, W., Schade, P. F., Hartmann, T. S., Kreiman, G., & Livingstone, M. S. (2019). Evolving images for visual neurons using a deep generative network reveals coding principles and neuronal preferences. Cell, 177(4), 999–1009.

[35] Walker, E. Y., Sinz, F. H., Cobos, E., Muhammad, T., Froudarakis, E., Fahey, P. G., … & Tolias, A. S. (2019). Inception loops discover what excites neurons most using deep predictive models. Nature neuroscience, 22(12), 2060–2065.

[36] DiMattina, C., & Zhang, K. (2011). Active data collection for efficient estimation and comparison of nonlinear neural models. Neural computation, 23(9), 2242–2288.

